# Development and application of SNP markers to facilitate DUS testing in tomato

**DOI:** 10.64898/2026.06.16.732596

**Authors:** Mathilde Causse, Fredérique Bitton, Patricia Faivre Rampant, Renaud Duboscq, Aurélie Bérard, Chrystelle Jouy, Chiara Delogu, Hedwich Teunissen, Cecile Collonier, Isabelle Le Clainche, Damien Hinsinger

**Author notes:** CropXR Institute, Utrecht University, Padualaan 8, 3584 CH Utrecht, the Netherlands.

## Abstract

Variety registration in Europe requires the evaluation of Distinction, Uniformity, and Stability (DUS) based on multi-environment trials and extensive phenotyping. The integration of molecular markers into DUS testing offers opportunities to increase efficiency and reduce costs, particularly in tomato (*Solanum lycopersicum* L.), a species characterized by rapid varietal turnover. We developed a high-density SNP genotyping resource targeting gene-rich regions across the tomato genome and applied it to a panel of 300 varieties registered over the past five decades. Temporal patterns of genetic diversity were assessed and compared with those observed in a collection of heirloom accessions predating 1970. Genome-wide association studies (GWAS) were conducted for 50 DUS traits to identify marker-trait associations and evaluate the potential of SNPs to complement phenotypic descriptors. Detected associations were compared with previously reported genes and quantitative trait loci (QTLs), enabling the validation of known loci and the identification of novel candidate genomic regions underlying trait variation. Finally, we assessed the discriminatory power of selected subsets of informative SNPs for variety distinction and grouping. Our results demonstrate the potential of integrating genomic and phenotypic data to enhance the robustness, resolution, and scalability of DUS testing in tomato.

## Introduction

Tomato breeding represents one of the most dynamic sectors in horticultural crop improvement, with more than 300 new varieties registered annually in Europe. To date, nearly 9,900 tomato varieties are listed in the EU Plant Variety Portal (https://ec.europa.eu/food/plant-variety-portal), reflecting both the intensity of breeding efforts and the high level of phenotypic diversification achieved in this species. In Europe, both the registration for marketing purposes (National Listing) and the granting of Plant Breeders’ Rights require to assess the Distinctness, Uniformity and Stability (DUS) of the varieties following harmonized technical protocols defined by the Community Plant Variety Office (CPVO). For vegetable species, DUS tests require candidate varieties to be compared with reference collections over at least two growing seasons, under conditions representative of their intended cultivation environments (open field, protected cultivation, or heated greenhouse), and a large number of phenotypic traits are characterized. For tomato, Distinction testing relies on the evaluation of 67 standardized descriptors assessed by official examination offices (EOs) across multiple countries (CPVO, 2025). These descriptors encompass plant architecture, floral traits, fruit morphology, organ colors, as well as resistance to a range of pathogens, sometimes considering multiple pathogen races. Traits are predominantly recorded as qualitative variables, including binary classes (e.g., presence/absence or resistant/susceptible) and ordinal scales, often ranging from 1 to 9, corresponding to continuous phenotypic variation. For fruit color and shape, each class corresponds to a specific state of expression of the characteristic. The list of descriptors has evolved over time, notably through the integration of new disease resistance traits such as those targeting Tomato Yellow Leaf Curl Virus (TYLCV), reflecting ongoing breeding priorities.

Beyond registered varieties, tomato genetic resources are exceptionally rich at the global scale, with more than 60,000 heirloom varieties conserved in germplasm collections alongside their wild relatives (Ross, 1998). These collections, including those maintained by institutions such as INRAE (Salinier et al., 2022), preserve thousands of accessions characterized using standardized descriptors (International Plant Genetic Resources Institute (IPGRI), 1996), often overlapping with DUS criteria.

Despite the extensive phenotypic diversity of tomato varieties (Pons et al., 2023), molecular studies have consistently shown that cultivated tomato exhibits relatively low genetic diversity compared to its wild relatives and to many other crop species (Blanca et al., 2015). This apparent paradox reflects both the domestication bottleneck and subsequent breeding practices. Historical analyses of tomato diversity, such as those conducted by Schouten et al. (2019), indicate that genetic diversity declined from domestication through early European dissemination, but began to increase again from the 1980s onwards. This resurgence is largely attributed to the introgression of resistance genes from wild species and the diversification of market classes, particularly the expansion of cherry tomato types.

In parallel, tomato has emerged as a model species for plant genetics, with numerous genes and quantitative trait loci (QTLs) underlying key agronomic traits having been identified and cloned (Rothan, Diouf et Causse, 2019). Notable examples include the *sp* gene controlling determinate growth habit (Pnueli et al., 1998) and the *u* mutation affecting fruit green shoulder (Powell et al., 2012a), as well as major QTLs governing fruit size and shape -fw2.2 (Frary et al., 2000), fw3.2 (Chakrabarti et al., 2013), lc (Muños et al., 2011), fas (Cong, Barrero et Tanksley, 2008). In addition, at least 26 disease resistance genes have been characterized, some of which directly corresponding to DUS descriptors (Foolad et Panthee, 2012).

Advances in molecular genetics have transformed the analysis of tomato diversity, shifting from biparental QTL mapping approaches to genome-wide association studies (GWAS), enabled by the availability of a reference genome (The Tomato Genome Consortium, 2012) and high-density single nucleotide polymorphism (SNP) datasets (Sauvage et al., 2014). Early SNP arrays, such as those developed within the SolCAP initiative (Sim et al., 2012), have progressively been replaced by whole-genome resequencing approaches, providing unprecedented resolution for genetic analyses (Zhao et al., 2019).

The increasing number of candidate varieties has made DUS testing progressively more resource-intensive, due to the expansion of reference collections required for reliable distinction. At the same time, the cost of genotyping has decreased substantially, while SNP markers offer high reproducibility compared to earlier marker systems. These trends have stimulated growing interest in the integration of molecular data into variety testing (Tang, 2022; Zhang, 2023; Mendler-Drienyovszki et al., 2024), including the prediction of phenotypic traits and the genetic characterization of candidate varieties. Large collaborative initiatives such as the INVITE European project have specifically aimed to enhance the efficiency of variety testing procedures and to evaluate the potential role of molecular markers across multiple crop species, including tomato (https://www.h2020-invite.eu/). In this context, the integration of high-throughput molecular tools into variety testing frameworks offers promising opportunities to complement conventional DUS evaluations.

Here, we developed a novel SNP genotyping resource based on 20,000 SPET (Single Primer Enrichment Technology) markers, strategically designed to target a large set of genes distributed across the tomato genome. This marker panel was conceived to maximize both genome coverage and functional relevance, with the objective of capturing genetic variation underlying key agronomic and DUS-related traits. Using this SNP panel, we genotyped a collection of 300 tomato varieties provided by several official examination offices (EOs), representing a broad spectrum of contemporary breeding material. These varieties were characterized at both molecular and phenotypic (DUS traits) levels, enabling a comprehensive assessment of their genetic diversity and temporal evolution, in order to better understand how recent breeding practices have shaped the genetic structure of cultivated tomato. We also compared this diversity to the diversity present in a collection of heirloom varieties gathered before 1970, genotyped with 7700 SNPS and characterized for several DUS traits.

To link genotype and phenotype, we conducted genome-wide association studies (GWAS) for a range of DUS traits, and compared the identified associations with previously characterized genes and QTLs, when available. This approach allowed us to validate known genetic determinants as well as to identify novel candidate regions potentially involved in trait variation. Several significant associations were also detected in the collection of genetic resources. Finally, we explored the potential application of molecular markers for variety distinction by designing small subsets of highly informative SNPs. These marker sets were optimized for their discriminating power among varieties, with the aim of providing practical tools to support or partially replace phenotypic assessments in DUS testing following validated UPOV models (UPOV, 2020).

## Materials and Methods

### Plant Materials and phenotypes

We gathered seeds of 300 varieties from three EOs, NAKT (Naktuinbouw, in the Netherland), GEVES (Groupe d’Etude et de contrôle des Variétés Et des Semences, France) and CREA (Council for Agricultural Research and Economics, Italy). Phenotypes for 67 traits (**Supp Table 1**) used for the variety registration were also provided by EOs with the date of variety registration. All the varieties were coded by CPVO as their data were produced using reference samples provided for official DUS purposes and remain confidential to preserve breeders’ interest. A collection of 272 heirloom varieties (named GR collection) was provided by CRB-Lég of INRAE UR-GAFL (Salinier et al, 2022).

### SNP design and genotyping

DNA was extracted using the “DNeasy Plant Mini Kit” (Qiagen Sciences, Germantown, MD, USA) following the manufacturer’s protocol, with RNase A (Qiagen) to remove any remaining RNA. The amount of DNA was quantified using a Qubit Fluorometer 3.0 and the “Qubit DNA Broad Range Assay Kit” (Invitrogen, Carlsbad, CA, USA). DNA purity (260/230 nm and 260/230 nm ratios) was assessed using the NanoDrop 1000 Spectrophotometer (Thermo Fisher, Waltham, MA, USA). Sample’s integrity was assessed on a 1% agarose gel.

We designed a new SNP panel : 20,000 SNPs were selected to tag as many genes as possible and represent SNPs previously used. The SNPs corresponded to close markers for 98 candidate genes and 27 disease resistance genes (Rothan et al, 2019), as well as SNPs from previous sets: (3640 SNPs from the SolCap array (Sim et al., 2012); 4287 from the CBSG array (Víquez-Zamora et al., 2013), 4182 from a previous SPET array (Barchi et al., 2019), and 1322 used in the MAGIC population (Pascual et al., 2015). A complement of 6441 SNPs was selected following a polymorphism screening in a set of 135 re-sequenced accessions including 53 *S. lycopersicum* (SL), 59 *S. l. cerasiforme* (SLC), 5 *S. pimpinellifolium* (SP) and 17 modern F1s, based on several criteria (minimum 5 reads, minimum MAF of 5%, no other variant (SNP or indel) in +/-50bp, at least one variant in each category (SL, SLC, SP and F1) and proximity to a candidate gene or QTL. A final set of 20,000 SNPs was selected to genotype the DUS collection using SPET Tecan technology. DNA concentrations were normalized based on fluorimetric quantification (Fluoroskan with Quant-it dsDNA - BR and HS kits, Thermo Fisher Scientific, Waltham, MA, USA). Libraries were built following Allegro Targeted Genotyping V2 protocol according to manufacturer (TECAN, Redwood City, CA, USA), with individual gDNA input ranging from 30 to 60ng. Libraries were then quantified by qPCR (KAPA Library Quantification Kit for Illumina Libraries - KapaBiosystems), and their profiles assessed on the Agilent Bioanalyzer using a High Sensitivity DNA kit (Agilent Technologies, Santa Clara, CA, USA). One pool showed remaining adapters dimers, thus requesting an additional purification step, performed with 0.8X v:v Ampure XP (Beckman Coulter, Brea, CA, USA). Libraries were pooled equimolarly and sequenced on a NovaSeq6000 instrument (Illumina, San Diego, CA, USA) in 100bp single-end mode, at the Genoscope, CEA—Institut de biologie François Jacob (Evry, France). Demultiplexing was performed using the eight first bases of the read index 1 as individual barcodes, with the next 6 bases used as UMI and written in a separate fastq file ; trimming was done as described in (Alberti et al., 2017), with an additional step using cutadapt (Martin, 2011) to remove UMI reads containing N (“--max-n 0”).

In addition to the target SNPs, 16,000 extra SNPs located in close proximity (within 50 bp) to the target SNPs were also detected. As these additional SNPs were in strong linkage disequilibrium with the target SNPs, only the target SNPs were retained for further analysis. After filtering for missing data (<15%) and MAF (> 0.05), a total of 14,107 SNPs were selected for further analyses. The GR collection had been earlier genotyped with 7650 SNP of the SolCAP array (Blanca et al. 2015) and phenotyped for 19 traits common to DUS characteristics (**supp table 2**).

### Statistical analysis

After removing duplicate individuals and those with more than 30% missing data, the analyses were performed on 281 DUS varieties and 272 GR accessions. For each SNP, we assessed the allele frequencies, heterozygosity, missing data (NA) frequency, and minimum allele frequency (MAF). The polymorphism information content (PIC) was calculated for each SNP as 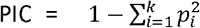 and averaged per chromosome on the whole set of varieties, and per group of registration decades.

The distribution of each trait was analyzed. Correlations between traits were calculated and a multivariate analysis was then performed on the DUS collection using R statistical software (R Core Team, 2021). GWAS was performed using the GAPIT R package (Wang et Zhang, 2021) including Kinship and Structure in the models. We compared the results of several models (MLM, FArmCPU, BLINK). We retained FarmCPU results as they were the most efficient to detect associations close to known genes. A Bonferroni multiple test threshold was used to determine significance. We also tested GMMAT R package (Chen et al., 2016), adapted to binary traits for GWAS analyses.

### Subset of markers

The discrimination ability of four SNP subsets was compared and we checked the number of similar varieties using either (1) 12 SNPs (one per chromosome) with highest PIC; (2) 12 SNPs selected among those with the most significant associations and related to a known gene; (3) 22 SNPs combining the two previous sets and (4) the 78 SNPs corresponding to the SNPs with at least one significant association.

## Results

### Phenotype diversity

The 67 DUS traits used to characterize the 281 accessions were evaluated using category traits with two to nine classes, based either on continuous scales or categorical variables. Their distributions are presented in **Supp Table 1**. Among the traits, eight phenotypic characteristics and all resistance traits (except nematode resistance) were coded as binary variables. Some characteristics were rarely observed, resulting in classes with very few individuals. This was particularly the case for absence of anthocyanin coloration on the hypocotyl of seedling, green stripes on fruit, orange flowers and the pinnate leaf type of blade (**supp table 1**). For 11 additional traits (including ten related to pathogen resistance), more than 70

% of the data were missing. Consequently, 50 traits were retained for subsequent analysis: four related to plant architecture, nine to leaf traits, three to inflorescences, 18 to fruit characteristics and 16 resistance traits corresponding to eight diseases (**Table 1**). GR accessions were characterized for 19 traits common to DUS descriptors (**Table 1 and supp table 2**). As they correspond to varieties cultivated before 1970, they carry very few disease resistances and were not characterized for these traits.

**Table 1:**
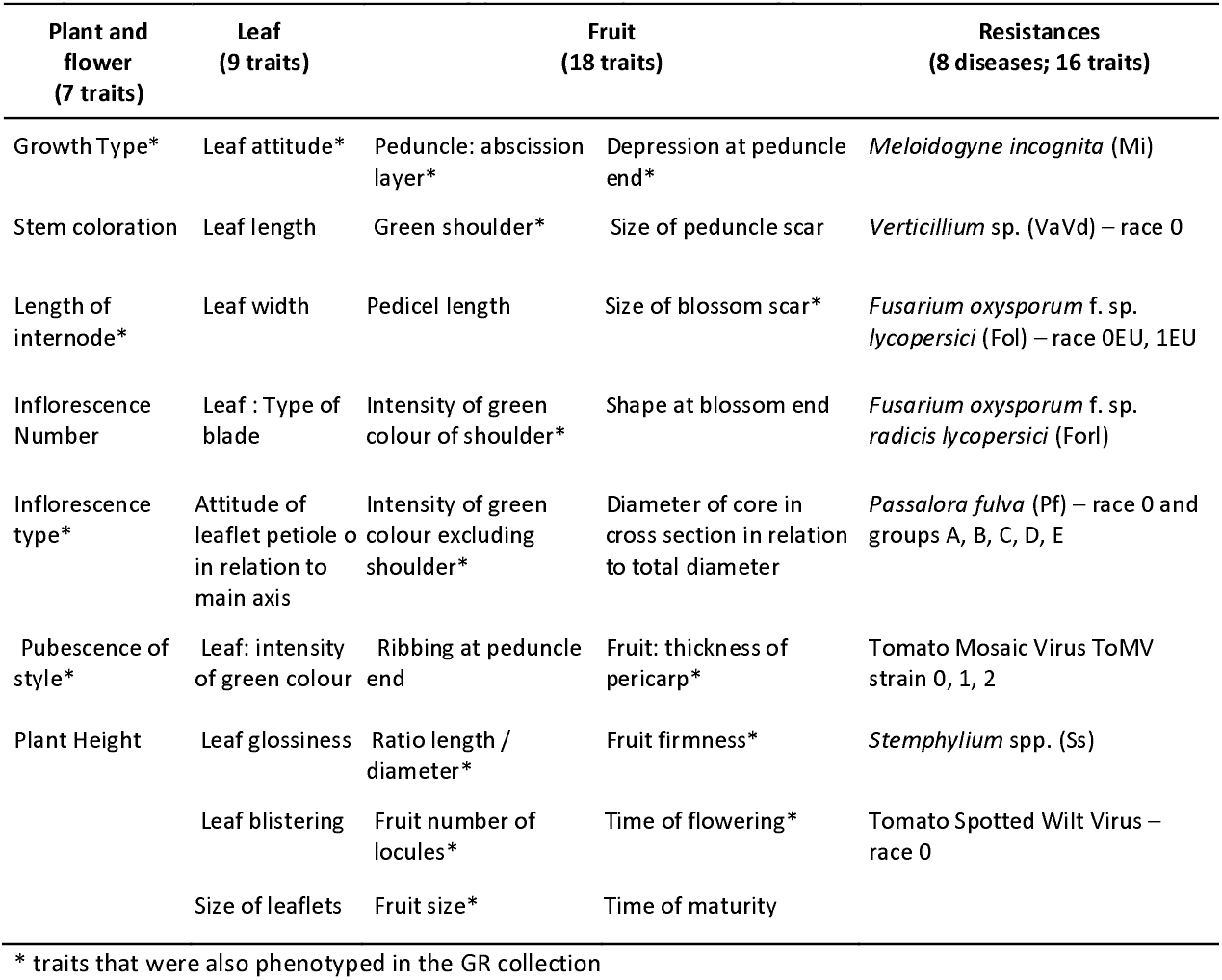
Traits used for analysis of the DUS panel (only those with sufficient data are shown). The complete list of DUS traits and their coding procedure is provided in **supp tab 1**

The varieties exhibited substantial phenotypic diversity. **Correlation analysis (supp figure 1)** revealed strong correlations among several groups of related traits, such as leaf length and width, or among fruit traits (locule number with blossom scar size, depression of peduncle, peduncle ribbing, fruit size and diameter). Some correlations reflect the use of different descriptors for the same trait (fruit shape and fruit length to diameter ratio) or closely related traits (time of flowering and maturity). Some disease resistances corresponding to different strains of the same pathogen were also correlated, notably for resistance to *Passafora fulva* (race 0 and groups A, B, C, D E) and Tomato Mosaic virus (ToMV, strain 0 and 1/2).

Overall diversity is illustrated by multivariate analysis (**Supp Figure 2**), with the first two principal components explaining 32 % of the variation. The first axis primarily separates varieties according to fruit size, and the second is associated with the presence or absence of green shoulder and resistances to *Verticillium* and *Fusarium oxysporum*. When growth type is projected onto the PCA, the lower diversity of determinate varieties is apparent (**Supp figure 3**).

Several traits have evolved over time (**supp table 3**). A major change observed since the 1970’s is the increasing prevalence of the absence of green shoulder: this trait was present in only 25 % of the GR accessions, but in 60% in the DUS panel. A similar trend was observed for inflorescence type, with only 32% of the GR accessions exhibiting predominantly uniparous inflorescences compared to 81% of the varieties in the DUS panel. The absence of a pedicel abscission layer, mainly used in processing tomatoes, emerged after 1990. Fruit size diversification also increased after the 2000’s, particularly with the registration of several cherry tomato varieties, and this diversification is also reflected in locule number and other fruit characteristics. Disease resistance genes have accumulated over time, with the average number per variety increasing from 0.60-1.00 in the 1970s-1980s to more than 4.00 in more recent varieties. Some varieties registered after 2020 carried up to 10 resistance genes (**Figure 1**).

**Figure 1:**
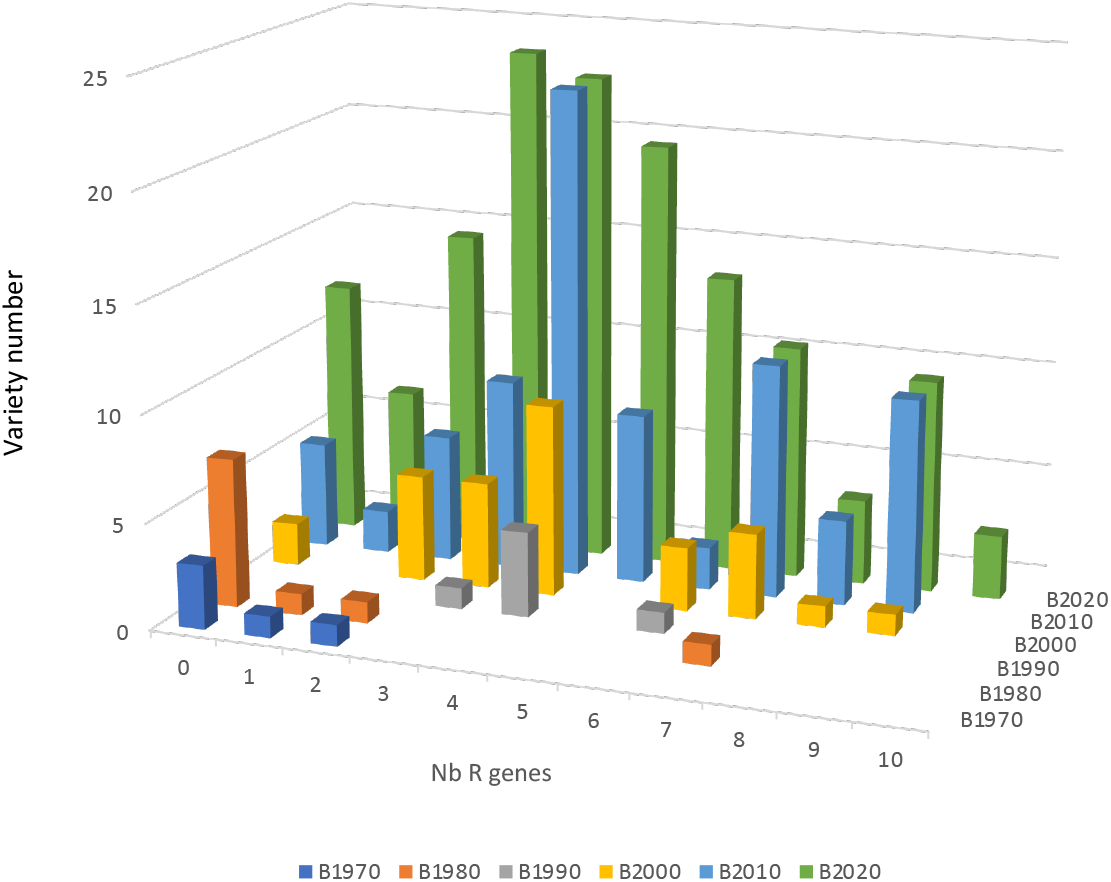
Number of varieties with 0 to 10 resistance genes per variety according to the registration period for 13 disease resistance traits

### Genotype diversity

Figure 2. shows the distribution on the 12 chromosomes of 14,107 SNPs used for analyses. A good coverage of the genome was obtained, with the number of SNPs per chromosome ranging from 402 on chromosome 8 to 2556 on chromosome 9. **Figure 3** illustrates the evolution of overall genotypic parameters across four time periods (with years prior to the 1990s grouped together), both genome-wide and per chromosome. The level of heterozygosity increased over time. A high proportion of homozygous varieties was observed before the 1980s, and an average heterozygosity of 2% of in the GR collection, consistent with their origin as self-pollinated varieties. Heterozygosity then increased steadily, from 12% before the 1990s to 26% after 2010. The polymorphism information content (PIC) followed a similar trend. The lowest values were observed before 1990 (mean of 0.17), with substantial variation among chromosomes during this first period. This was followed by a regular increase, reaching an average of 0.37 after 2010 (**Figure 3** and **supp table 4**).The proportion of missing data also increased over time, likely due to the introgression of disease resistance genes from wild relatives, whose sequences are not enough similar to the target sequences. In the GR collection the average PIC was 0.24 but with a limited variation between chromosomes, and MAF ranged from 0.08 to 0.18 across chromosomes. The genetic structure of this collection has been previously described in Blanca et al (2015).

**Fig 2:**
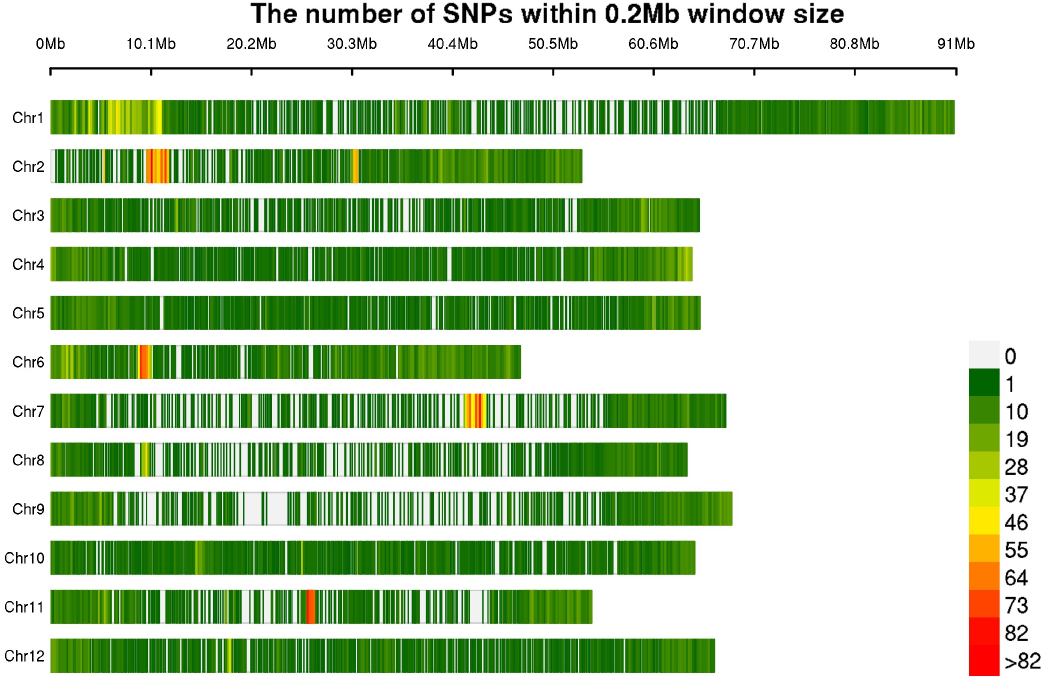
Distribution of the 14,000 SNPs used to genotype the DUS panel on the 12 chromosomes

**Figure 3:**
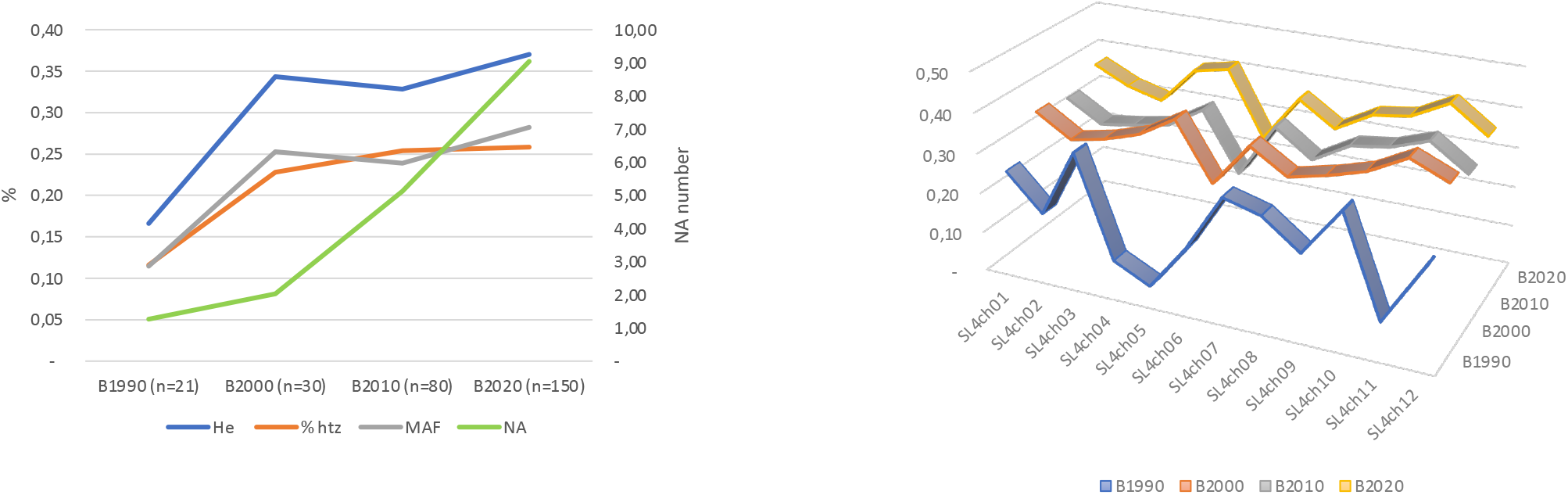
Genotypic parameters over the registration periods: a. Evolution of PIC (He), observed heterozygosity (% htz) and MAF over the periods (in % on the left axis); on the right axis, average number of missing data per accession. b. He values per chromosome and period of registration

### GWAS for DUS traits

We performed GWAS with the DUS traits in order to identify new markers to accelerate distinction trials and check whether known genes could be helpful in the process. For the GWAS, several models implemented in the GAPIT package (GLM, MLM, FarmCPU, BLINK) were evaluated. The results obtained with the FarmCPU model (Chen et al., 2016) are presented here, as this approach provided the most consistent outcomes, both in terms of QQ-plots and of detection of associations with previously characterized genes. The GMMAT package, specifically designed for binary traits, was also tested and yielded largely similar results with the most significant associations and a few additional ones. When genes known to control a given trait were available, we examined whether they co-localized with significant associations. In total, 97 associations were detected for 36 traits, with one to six associations per trait, while no significant association was detected for 14 traits. The detailed results are provided in **supp table 5. Figure 4** presents the genomic positions of significant associations and the corresponding genes when co-localization was observed.

**Fig 4:**
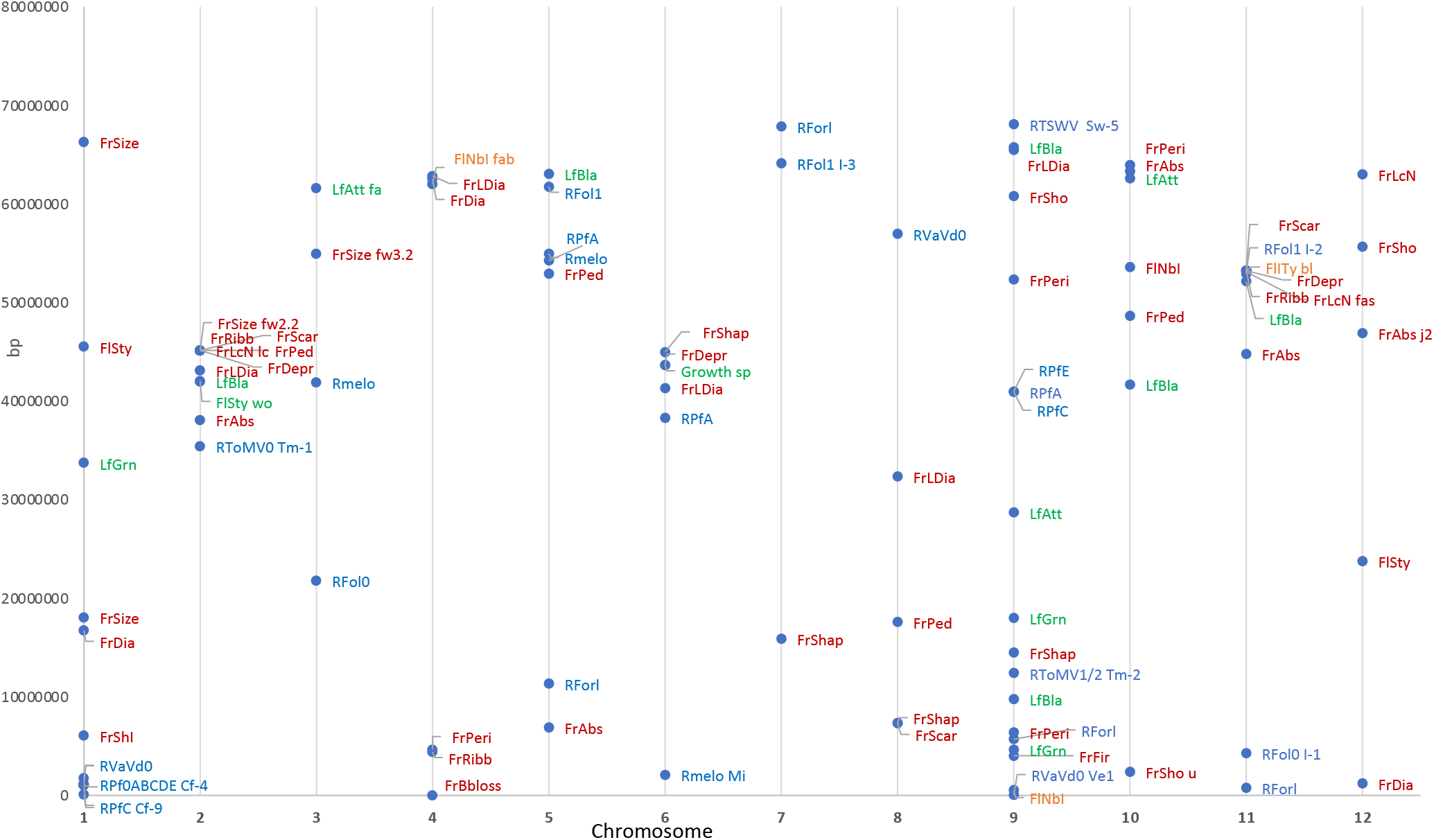
Map of the associations detected in the DUS panel by GWAS **Code of traits :** Growth Type - Growth ; Leaf Attitude - LfAtt; Leaf Blade Type - LfBla; Leaf Green Intensity - LfGrn; Style Pubescence - FlDty; Nb Inflorescence - FlNb; Peduncle Abscission Layer - FrAbs; Inflorescence Type - FlITy; Green Shoulder - FrSho; Green Shoulder Intensity - ; Locule Nb - ; Fruit Blossom Scar Size - ; Fruit Depression Peduncle - FrDepr; Fruit Ribbing Peduncle - FrRibb; Fruit Peduncle Scar Size - FrScar; Fruit Size - FrSize; Fruit Diameter - FrDia; Fruit Length To Diam -FrLDia; Fruit Shape Longitudinal - FrShap; Fruit Firmness - FrFir; Fruit Pericarp Thickness - FrPeri; Fruit Shape Blossom End - FrBloss; Resistance Fol0EU – Rfol0; Resistance Fol1EU – Rfol1; Resistance Forl - RForl; Resistance Mi - Rmelo; Resistance Pf0 – RPf0; Resistance PfA - RPfA; Resistance PfB - RPfB; Resistance PfC - RPfC; Resistance PfD - RPfD; Resistance PfE - RPfE; Resistance ToMV0 – RtoMV0; Resistance ToMV1/2 – RtoMV1/2; Resistance VaVd0 – Rvavd0; Resistance TSWV – RTSWV

For Growth Type, a highly significant association (P-value =7.10^-46^) was identified on chromosome 6, corresponding to the *sp* gene (Pnueli et al., 1998). Classification accuracy was high, with only six out of 281 varieties incorrectly classified. For Leaf attitude, three QTLs were detected on chromosomes 3, 9 and 10; however, none were near the *Erl* gene, recently shown to control the semi-erect phenotype (Fernandes et al., 2025). For leaf blade type, six QTLs were detected. Although several mutations in different genes have been identified (*entire, lanceolate, wiry*; (Hareven et al., 1996), none co-localized with the QTLs detected. For leaf green intensity, three QTLs were detected. Regarding flower traits, one of the three QTL associated with style pubescence was located 400 kb from the gene underlying the *Wooly* mutation (Yang et al., 2011).

For the number of inflorescences, three QTLs were detected, but none near the *s* gene, which confers compound inflorescence (Lippman et al., 2008). For inflorescence type, a single association was detected, located 1.5 Mb from the *blind* locus on chromosome 11.

For Peduncle Abscission Layer presence, two genes, *j* and *j-2*, have been previously described (Mao et al., 2000; Budiman et al., 2004). Among the five associations detected, one could correspond to the *j-2* locus. For Green shoulder, one highly significant association (P-value =3.10^-26^) was detected on chromosome 10, located very close (234 bp far) to the *uniform color (u)* mutation, which is encoded by a Golden 2-like transcription factor (Powell et al., 2012a). A single association for green shoulder intensity was detected on chromosome 1.

For Locule number, three associations were detected, two of them were located near the genes corresponding to *lc* (a mutation in the regulatory region of the Wuschel gene, on chromosome 2; Muños et al., 2011) and *fas* (a YABBY-like transcription factor on chromosome 11; Cong, Barrero et Tanksley, 2008). When both genes were considered together, the number of locules of 87% of the varieties was accurately predicted (**supp table 6**). These two SNPs were also significant together with other QTLs for fruit blossom scar size (three QTLs), fruit depression of peduncle (three QTLs), fruit ribbing peduncle (three QTLs). The *lc* locus alone was significant among the four QTLs for fruit peduncle scar size. For Fruit size, four QTLs were detected: one near *Lc*, one on chromosome 3 at 4 Mb from fw3.2 (Chakrabarti et al., 2013) and two on chromosome 1. At least two distinct associations were detected for fruit diameter.

For fruit shape, which corresponds to two traits (Fruit Length to Diameter ratio and Fruit Shape Longitudinal), nine associations were detected. However, they were located relatively far from known genes (*ovate, Sun* or *fs8*.*1*; Van Der Knaap et al., 2014; Wu et al., 2018). One association on chromosome 4 was common to Fruit length to diameter ratio and fruit shape at the blossom end. Finally, for fruit firmness, one association was detected on chromosome 9, and four were identified for pericarp thickness. For disease resistances, 24 associations were detected, eight being located in the vicinity of known disease resistance genes (Rothan, Diouf et Causse, 2019). For Fusarium resistance, distinct associations corresponding to the three genes *I-1, I-2* and *I-3* were detected. For Fol0, the main association was 14 kb from *I-1* (with a P-value =3.10^-34^). For Fol1 three associations were detected, including a major one 50 kb from *I-2* gene and another 628 kb from *I-3* gene. For Forl four associations were detected, including a major locus on chromosome 9, where a gene for this traits has previously been mapped (Kim et al., 2016). For the nematode resistance, the *Mi* gene was detected among three associations, although not with a strong P-value. For resistance to *Passalora fulva* (Pf) (formerly *Fulvia fulva* (Ff) and *Cladosporium fulvum*, Cf), six traits were scored on only 72 to 116 varieties. Very few differences were observed between the scores for the different races. The *Cf-4* gene was consistently detected across all races, while *Cf-9*, located on top of chromosome 1, was associated with PfC. Another association on chromosome 9 was detected for the race PfC. For ToMV, the *Tm-1* gene was detected on chromosome 2 for the strain ToMV0 and a highly significant association was detected for the strains ToMV1/2, on chromosome 9, in the region of *Tm-2* gene. For Verticilium, three associations were detected, with the major one located at 7 kb from the *Ve* gene. Finally resistance to TSWV was characterized only on 103 varieties, but a single association was detected 7 kb from *Sw5*.

The GMMAT package designed for binary traits yielded results consistent with those obtained using FarmCPU for the most significant associations. Out of 49 significant associations, 39 were also detected with FarmCPU and 10 were uniquely detected by GMMAT (**supp table 5**). We further analyzed the fruit and flesh color with this package by testing specific contrasts (red vs each alternative color). This analysis revealed associations near the mutation tangerine (t) for orange color, y for pink epidermis, r for yellow fruit and hp-3 for the brown color and green flesh (**supp table 7**).

No significant association (P<10^-5^) was detected for 14 traits. These included traits with unbalanced distributions (leaf blistering, leaf glossiness, fruit extent of green shoulder), but also size-related traits (leaf length and width, peduncle length, leaflet size, plant height) and developmental traits (time of flowering and maturity). We tested genomic prediction for plant height, flowering time, and maturity, and showed strong correlations between predicted and observed phenotypes (**Supplementary Figure 5**), suggesting the involvement of numerous small-effect QTLs, but a high predictability of the model.

The Genetic Resources (GR) collection was phenotyped for 19 traits common to DUS descriptors. A total of 60 associations were detected for 14 traits (**supp table 8**). Among these, 14 overlapped with associations detected in the DUS panel, and 11 corresponded to known genes, including *u, sp, Lc, fas, fw3*.*2, fw11*.*2, fw6*.*1, j-2* and *r*.

### Discrimination of the varieties

When a major association was identified, the proportion of variety correctly classified based on their genotype at the peak-SNP was evaluated (**Table 2**). Among 19 associations across 17 traits, seven genes showed classification accuracies above 90%. For four genes, accuracy exceeded 80%, for five genes, it was above 70% and for three genes it was below 70%.

**Table 2:** Discrimination power of SNP peaks close to known genes.

| Trait | Chromosome | Position of closest SNP in GWAS | % well classified | Candidate gene | Gene code | SL4.0 Position | Distance SNP peak-gene (kb) |
| --- | --- | --- | --- | --- | --- | --- | --- |
| Growth type | SL4ch06 | 43636571 | 98 % | sp | Solyc06g074350 | 43636031 | 0.540 |
| Green shoulder | SL4ch10 | 2403680 | 83 % | u | Solyc10g008160 | 2169302 | 234.378 |
| Locule number | SL4ch02 | 45149080 | 86 % | lc | Solyc02g083950 | 45191157 | 42.077 |
| " " | SL4ch11 | 53127617 | 51% | fas | Solyc11g071810 | 53232733 | 105.116 |
| Fruit size | SL4ch02 | 45149080 | 74 % | fw2.2 | Solyc02g090730 | 50290300 | 5141.220 |
| " " | SL4ch03 | 54936943 | 51 % | fw3.2 | Solyc03g114940 | 58848000 | 3911.057 |
| Inflorescence nb | SL4ch04 | 62012066 | 77 % | fab | Solyc04g081590 | 63504450 | 1492.384 |
| Leaf attitude | SL4ch03 | 61590999 | 73 % | ? | ? |  |  |
| Peduncle abscission | SL4ch12 | 46891007 | 94% | j-2 | Solyc12g038510 | 50050180 | 3159.173 |
| Resistance ToMV0 | SL4ch02 | 35415571 | 69 % | Tm-1 | Solyc02g062560 | 32290560 | 3125.001 |
| Resistance ToMV1/2 | SL4ch09 | 12400716 | 97 % | Tm2 | Solyc09g018220 | 13656895 | 1256.179 |
| Resistance Mi | SL4ch06 | 2063628 | 89 % | Mi-1 | Solyc06g008410 | 2352904 | 289.276 |
| Resistance VaVd0 | SL4ch09 | 51343 | 95 % | Ve-1 | Solyc09g005090 | 58717 | 7.374 |
| Resistance Fol0EU | SL4ch11 | 4263221 | 98 % | l-1 | Solyc11g011180 | 4277933 | 14.712 |
| Resistance Fol1EU | SL4ch11 | 52928245 | 90 % | l-2 | Solyc11g071430 | 52978257 | 50.012 |
| Resistance Forl | SL4ch09 | 5706554 | 93 % | Fr1 | ? |  |  |
| Resistance Cf (race 0, B, D) | SL4ch01 | 1132996 | 85 % | Cf-4 | Solyc01g006550 | 1130793 | 2.203 |
| Résistance Cf (race C) | SL4ch01 | 81863 | 72 % | Cf-9 | Solyc01g005160 | 151978 | 70.115 |
| Résistance TSWV0 | SL4ch09 | 68091467 | 71 % | Sw-5 | Solyc09g098130 | 68083873 | 7.594 |

The primary objective of the DUS test is to demonstrate that a new variety is distinct from the existing ones. We thus tested the discriminatory power of four sets of SNP markers selected from those significantly associated with traits. The SNP lists and results are presented in **supp table 9** and **Figure 5**). To support distinctness testing and the selection of control varieties, a predefined set of grouping traits is used (**supp Table 1**). Using a set of 12 SNPs significantly associated with grouping traits and corresponding to candidate genes (i.e., those with lowest P-values), 58 out of 281 varieties were indistinguishable from at least one other variety. When using 12 SNPs selected as one per chromosome with the highest MAF, only eight pairs and one triplet of undistinguishable varieties were observed (6.7% undistinguished). Combining these two sets into a set of 22 SNPs reduced the number of indistinguishable varieties to only three pairs. Finally, when all 78 significant SNPs were considered, only a single pair of identical varieties remained. We further checked this pair for the full dataset of 14,000 SNPs and they differed for 300 SNPs (2%), among which 195 were on chromosome 9. In addition, the two varieties differed for 14 phenotypic traits.

**Figure 5:**
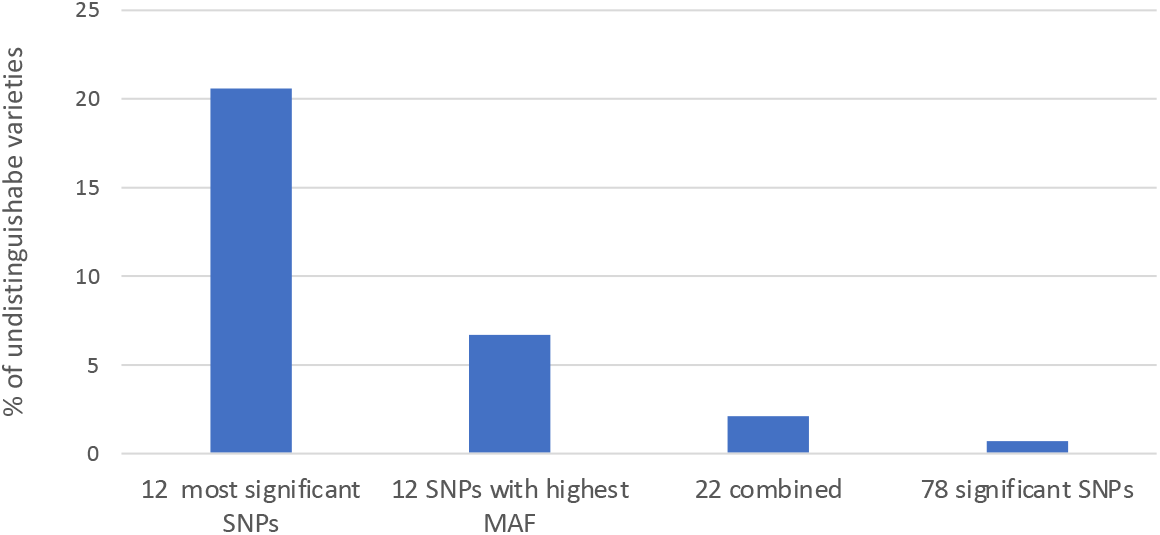
Discrimination power (percentage of undistinguishable varieties) of four SNP subsets. The composition of the four subsets is detailed in supplementary table 9.

## Discussion

In DUS testing, a large number of traits are routinely assessed, many of which are inherently correlated and may result in redundant effort. This redundancy is particularly pronounced for fruit-related traits, especially those associated with locule number. A smaller, more strategically selected set of descriptors may be sufficient to capture the relevant phenotypic variation. Integrating molecular markers with phenotypic evaluation could further enhance the efficiency and accuracy of variety discrimination. Image-based phenotyping approaches could further improve the scoring accuracy and throughput for these traits (Li et al., 2025). For resistance traits, several correlations were also observed, for instance between Pf resistance genes and for TMV resistance to two strains. However, some discrepancies were detected in the GWAS results, suggesting partially independent genetic control.

A SPET panel of SNPs was developed for this study, prioritizing markers located near genes or QTLs associated with phenotypic traits. This panel could be readily applied to other diversity studies or GWAS analyses, especially as genotyping costs continue to decrease with technological advances. Genotypic data revealed clear temporal changes in population structure. Prior to 1970, most varieties consisted of pure lines (as also observed in the genetic resource collection), whereas F1 hybrids have become predominant since the 1990s. This shift, combined with an increase in the number of resistance genes per variety (from none in heirloom varieties to up to ten in modern cultivars), has led to a progressive increase in genetic diversity, as reflected by PIC values. These findings are consistent with those of Schouten et al. (2016), who reported similar trends in a smaller collection. These introgressions have also resulted in a higher proportion of missing genotype data, largely due to interspecific introgressions. The diversification of market classes, particularly the registration of cherry tomatoes exhibiting greater variability than large-fruited types (Blanca et al., 2015; Pons et al., 2023), has further contributed to this pattern.

Introgressions often involve large genomic fragments; for example, the *Tm-2* locus corresponds to an introgression exceeding 50 Mb (Van Rengs et al., 2022). As most of the disease resistance genes are dominant, such segments are frequently maintained in the heterozygous state, as they may carry deleterious alleles linked to genetic load.

We also documented temporal trends in key phenotypic traits. For example, the green shoulder phenotype has progressively disappeared, despite its reported negative impact on sugar content (Powell et al., 2012b). Similarly, the widespread use of the *j-2* mutation in processing tomatoes illustrates breeding-driven trait evolution.

GWAS analyses based on DUS traits and genotypic data identified a large number of marker–trait associations. Major loci such as *lc* and *fas* showed strong effects on fruit size and shape traits (Olivieri et al., 2026). Many associations mapped close to 18 previously characterized genes with large phenotypic effects. For several traits, a major association was detected along with a limited number of modifier loci (e.g., *sp, u*, or certain disease resistance genes), while in other cases, secondary associations likely reflected linkage disequilibrium with the main causal variant. For instance, in the case of TMV resistance, three SNPs on chromosome 1 were identified in complete LD with the *Tm2* locus on chromosome 9.

For other traits, multiple genes controlling similar phenotypes may segregate at different frequencies, complicating their detection in GWAS. In addition, some traits recorded as continuous variables may actually reflect distinct underlying genetic mechanisms. For example, leaf attitude is scored along a gradient from erect to drooping, although these extreme phenotypes may be controlled by different genes. Similarly, fruit shape showed several associations that were distant from known loci such as *ovate, SUN*, or *fs8*.*1*, possibly because each morphological class corresponds to specific combinations of mutations rather than a true continuous distribution. Rodriguez et al. (2011) demonstrated that combinations of *lc, fas, ovate*, and *SUN* alleles can explain up to 95% of fruit shape variation in a diverse collection. While several major genes are already well characterized, the predictive value of associated SNPs must be confirmed across broader germplasm collections. Conversion of these markers into KASP assays could facilitate their routine application in breeding and variety registration.

Several challenges were encountered in QTL detection. First, missing data arising from evolving phenotyping protocols—particularly for newly introduced resistance traits—reduced analytical power. Second, the SNP closest to a candidate gene was not always the most informative, especially in regions affected by wild introgressions, which tend to increase missing data rates. Third, certain traits exhibited highly unbalanced class distributions, with rare phenotypes limiting statistical power. This was particularly problematic for fruit color, where each category had to be analyzed separately despite small sample sizes. These constraints are at odds with standard GWAS practices, such as filtering out SNPs with minor allele frequency (MAF) below 0.15.

In the case of disease resistance genes introgressed from wild species, alleles are often maintained in the heterozygous state, resulting in missing or underrepresented genotypic classes and reducing marker informativeness. Regarding statistical methods, GMMAT proved effective for categorical traits such as color, but less so for other trait types.

The limited success in mapping QTLs for quantitative traits likely reflects strong environmental effects, given that the phenotype dataset spans over 50 years, multiple environments, and varying scoring practices. Changes in control varieties and evaluation standards may have further contributed to phenotypic variability. Notably, for three quantitative traits—plant height, flowering time, and maturity— no significant associations were detected, despite their typically high heritability (Bauchet et al., 2017). However, genomic prediction analyses showed strong correlations between predicted and observed phenotypes, suggesting a genetic architecture involving numerous small-effect QTLs, potentially with environment-specific effects.

Comparisons between GWAS results obtained from the genetic resource (GR) collection and the DUS panel revealed only a limited proportion of shared associations. This likely reflects differences in genetic diversity, as the GR collection captures early domestication processes, whereas the DUS panel reflects more recent breeding efforts. Consequently, some alleles may be already fixed or segregating at different frequencies across panels

In terms of varietal discrimination, a relatively small number of markers may be sufficient for initial classification and assignment to testing groups. The inclusion of key genes controlling fruit color and newly deployed resistance genes could further enhance discriminatory power, although validation on larger and more diverse sample sets is required.

## Conclusion

Research to use molecular markers to distinguish plant varieties and assess essential derivation have started many years ago. Advances in marker technology, from early PCR-based methods to today’s SNP assays, have driven major gains in reproducibility and cost-efficiency. These developments make molecular approaches increasingly practical for variety registration. Our results demonstrate that even a small panel of carefully chosen SNPs can substantially improve discrimination between varieties. When these genetic markers will be integrated with high-throughput phenotyping, the traditional DUS framework should be accelerated. Together, marker-assisted and phenotype-driven strategies offer a clear pathway for modernizing DUS testing: improving precision, reducing time and expense, and strengthening legal certainty in registration decisions. Adoption of these tools by testing authorities and breeders will be central to realizing their full potential.

## Supplementary materials

**Supp Table 1 :** List of all traits used in DUS trials. Distribution of varieties across classes

**Supp Table 2 :** Distribution of varieties in the GR panel for 19 DUS traits

**Supp Table 3 :** Distribution of some traits according to the registration period

**Supp Table 4 :** Genotype parameters in the DUS collection

**Supp Table 5 :** GWAS results in the DUS panel

**Supp Table 6 :** Distribution of Locule Number according to the genotypes at two SNPs (SL4ch02_45149080 for lcn and SL4ch11_53127617 for fas locus)

**Supp Table 7 :** Fruit color associations with GMMAT

**Supp Table 8 :** GWAS results in the GR panel

**Supp Table 9 :** SNP lists for discrimination

**Supp Figure 1 : C**orrelation matrix of traits for DUS trials

**Supp Figure 2 :** PCA on phenotypic traits and clustering according to growth type and green shoulder

**Supp Figure 3 :** Diversity of phenotypes according to year and growth type

**Supp Figure 4 :** Distribution of He, MAF and missing data per chromosome according to registration period

**Supp Figure 5 :** Prediction and results GWAs for (a) Plant Height, (b) Flowering Time and (c) Maturity time

## Acknowledgements

We are grateful to CEA-IbFJ-Genoscope for providing sequencing, bioinformatics and storage facilities. We acknowledge the CRB-Lég of INRAE UR GAFL for providing genetic resources and to Hélène Burck who characterized all these varieties. Thanks to Rafael Feriche-Linares for his help in starting the data analysis.

## Author contributions

MC: conceptualization, data analyses, and writing the manuscript; RD, DM, PFV, AB, ILC: Molecular biology; FB bioinformatic data analysis; CJ, CD, HT, CC: organizing samples and data collection; MC: funding acquisition

## Data availability

The positions of the SNP of the SPET panel and Genotypes for a few accessions, together with data (phenotypes and genotypes) of GR panel are available on https://doi.org/10.57745/6YAKI6. Phenotypes and genotypes of DUS data may be available upon request to Naktuinbouw (contact person).

