## Supplementary material for "Development and application of SNP markers to facilitate DUS testing in tomato": Supp Figures: Invite MS supp figures June26.pptx

### Slide 1
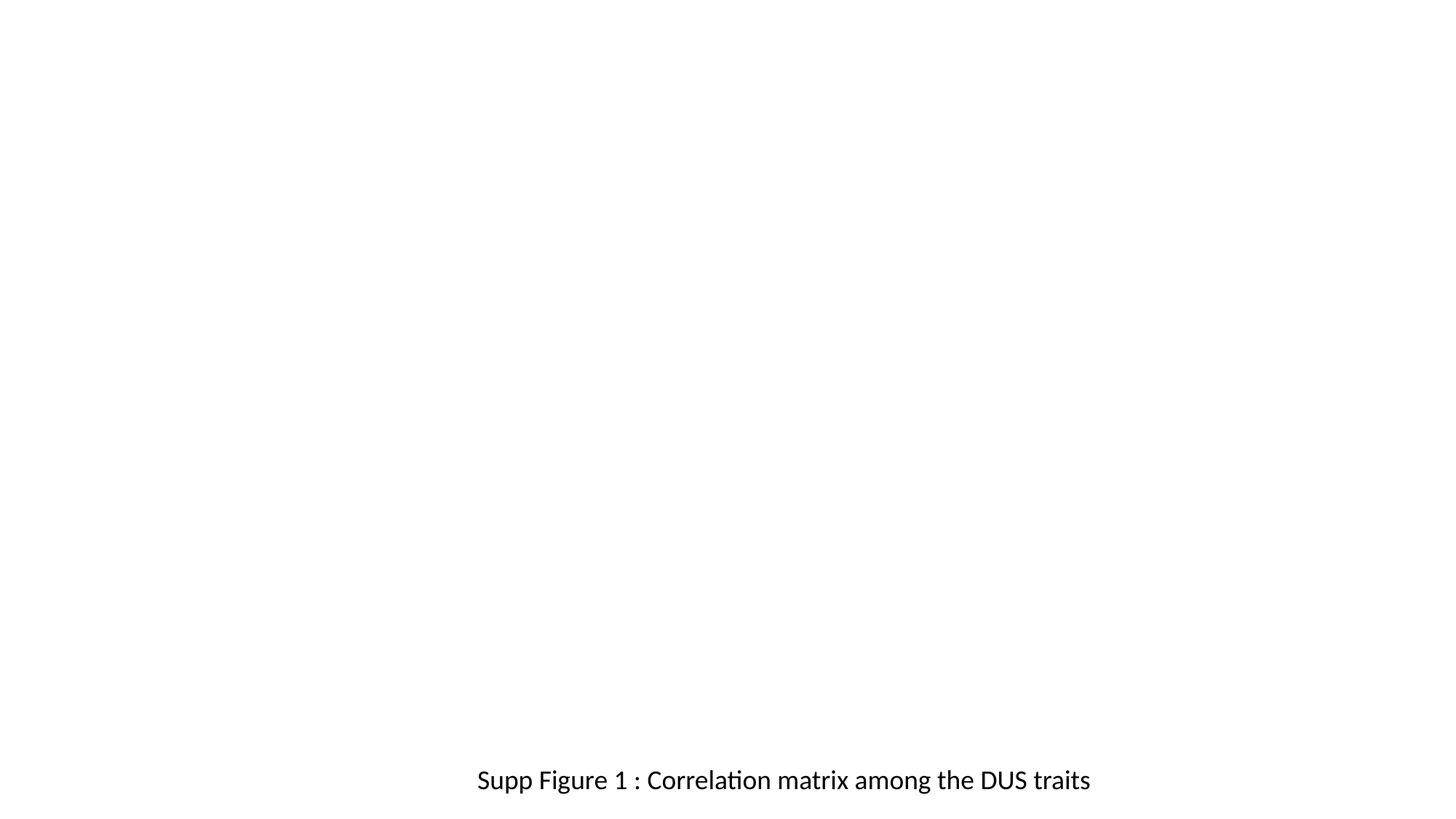

Supp Figure 1 : Correlation matrix among the DUS traits

### Slide 2
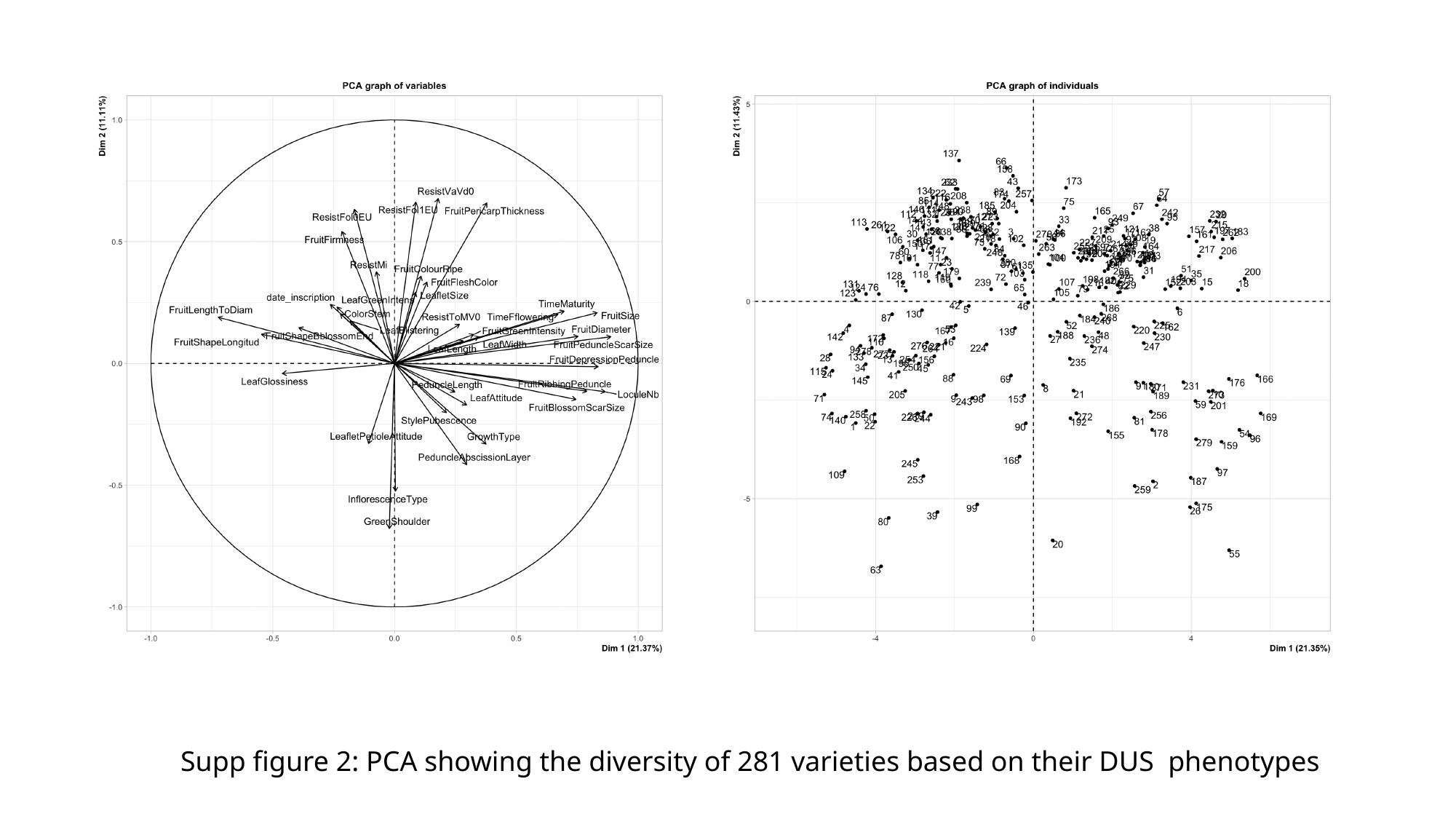

#### Supp figure 2: PCA showing the diversity of 281 varieties based on their DUS phenotypes

### Slide 3
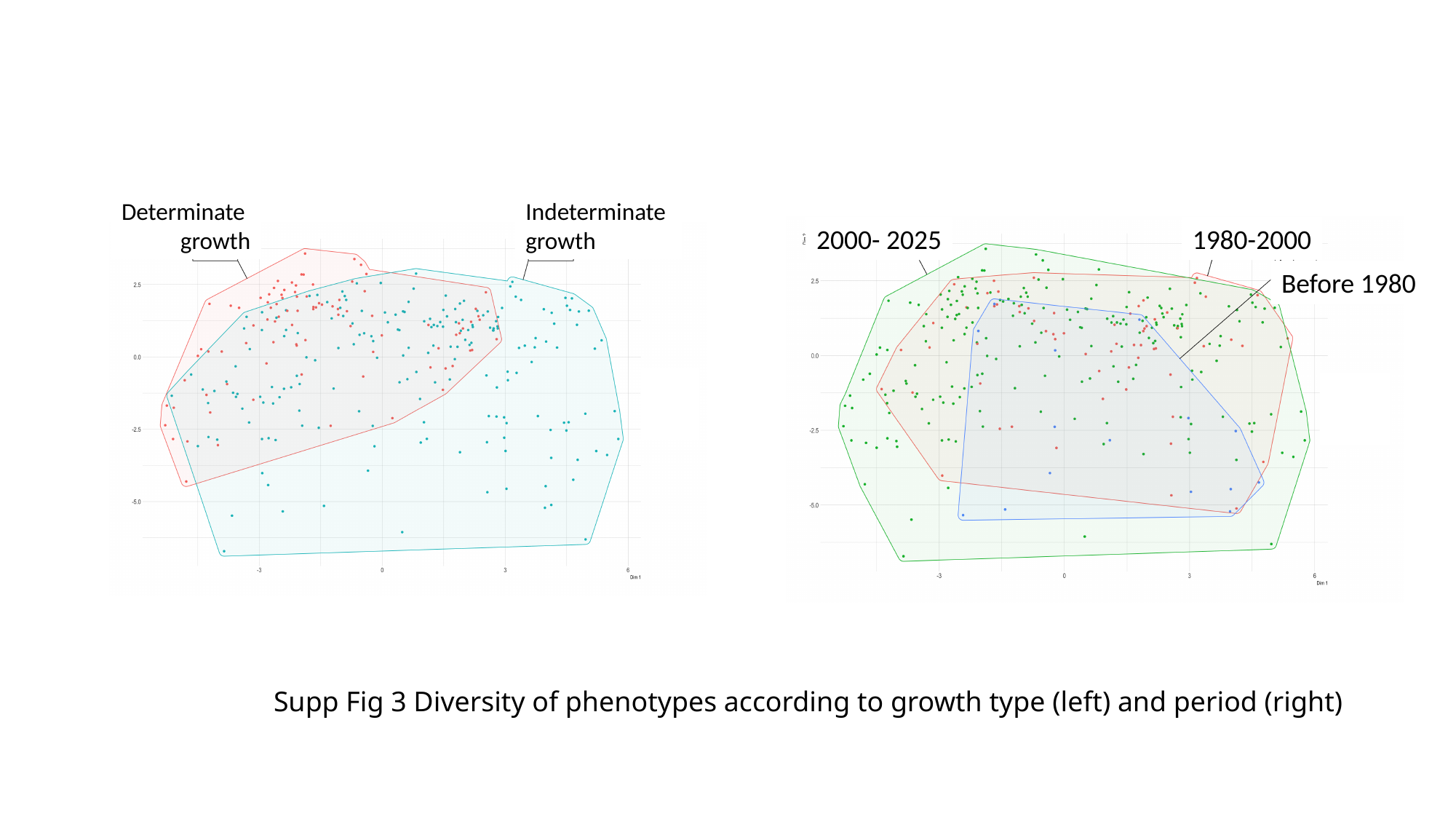

Determinate
growth
Indeterminate
growth
2000- 2025
1980-2000
Before 1980
#### Supp Fig 3 Diversity of phenotypes according to growth type (left) and period (right)

### Slide 4
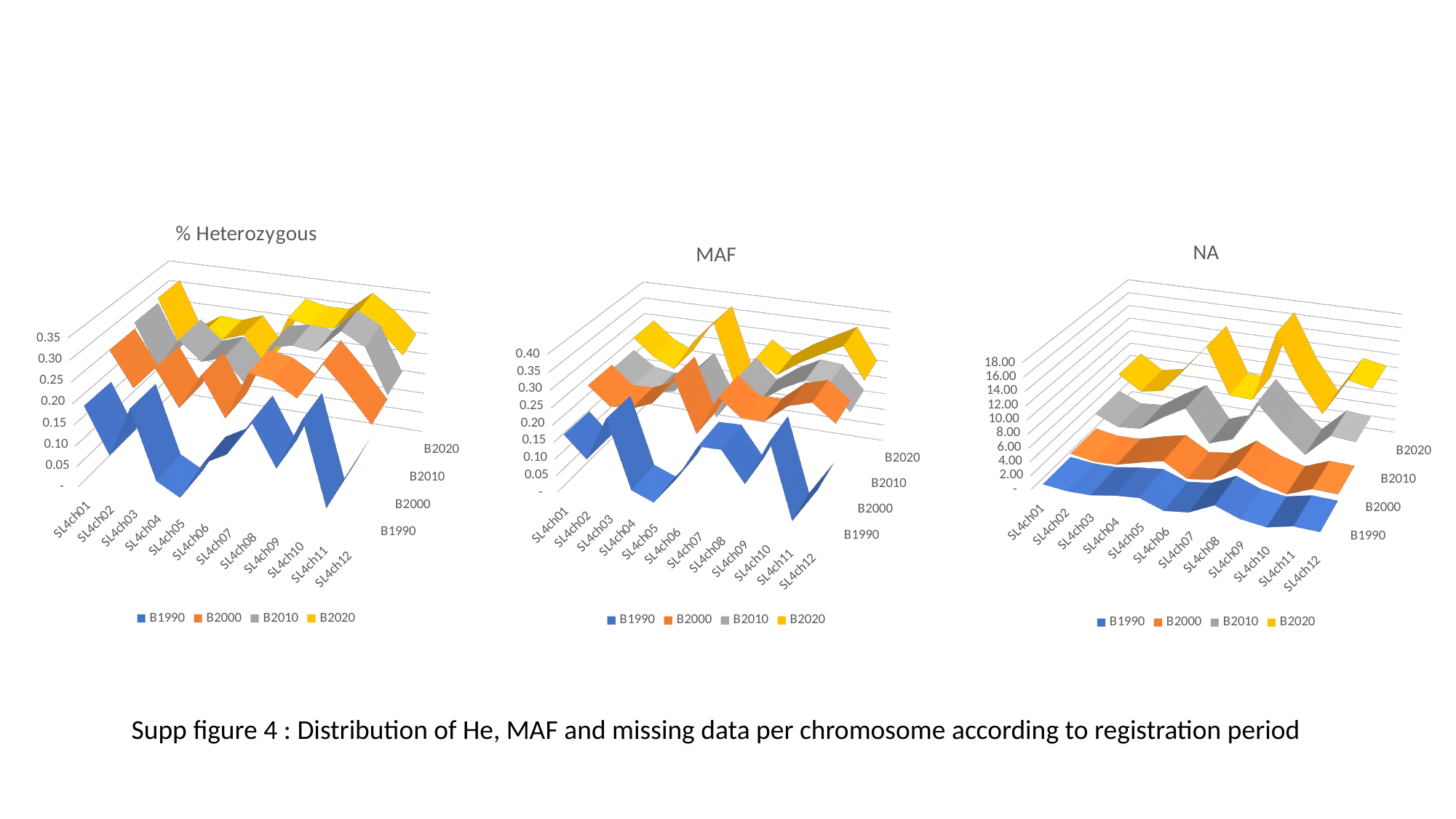

[unsupported chart]
[unsupported chart]
[unsupported chart]
Supp figure 4 : Distribution of He, MAF and missing data per chromosome according to registration period

### Slide 5
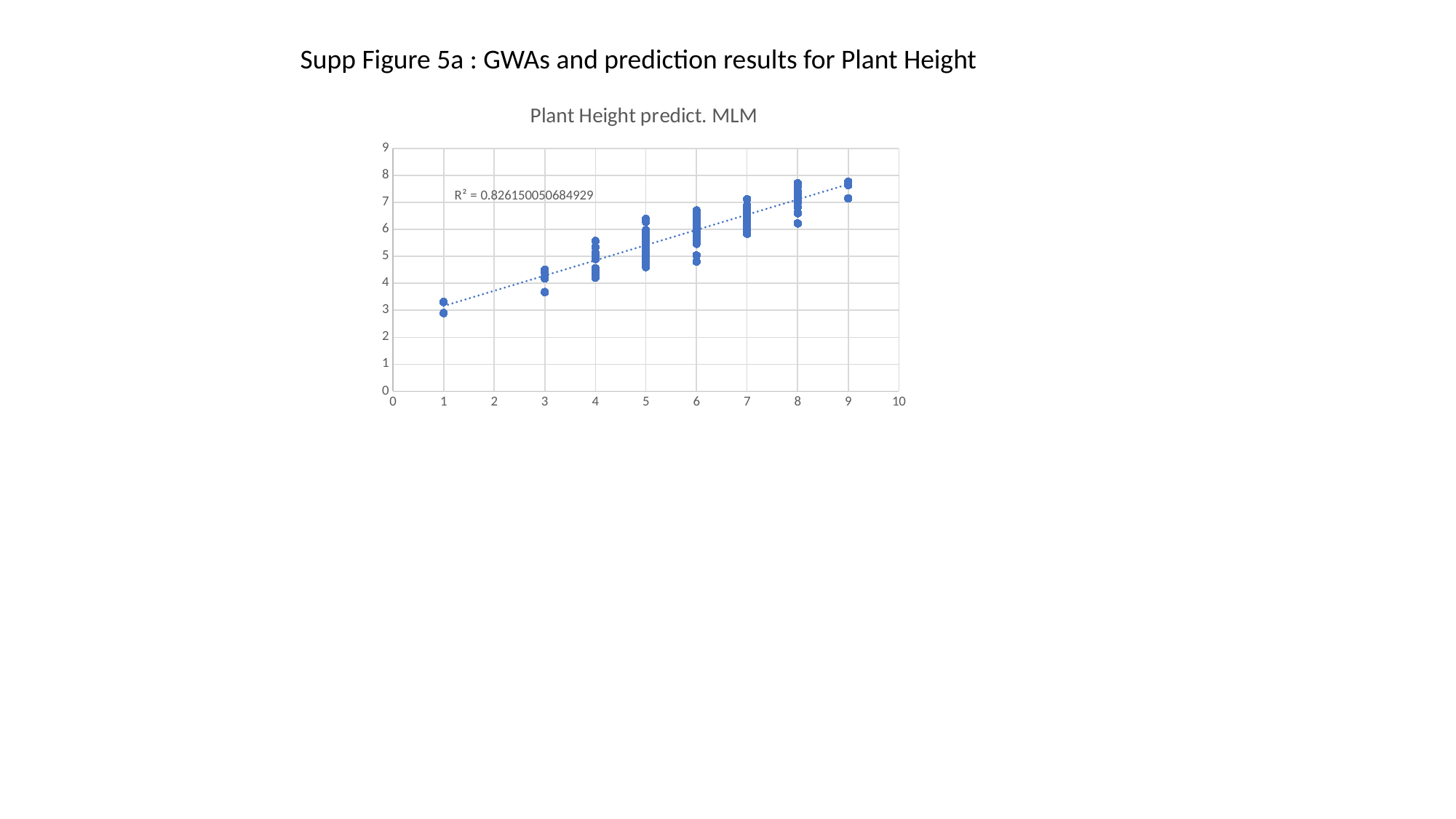

Supp Figure 5a : GWAs and prediction results for Plant Height
###### Chart: Plant Height predict. MLM
| Category | predict |
|---|---|

### Slide 6
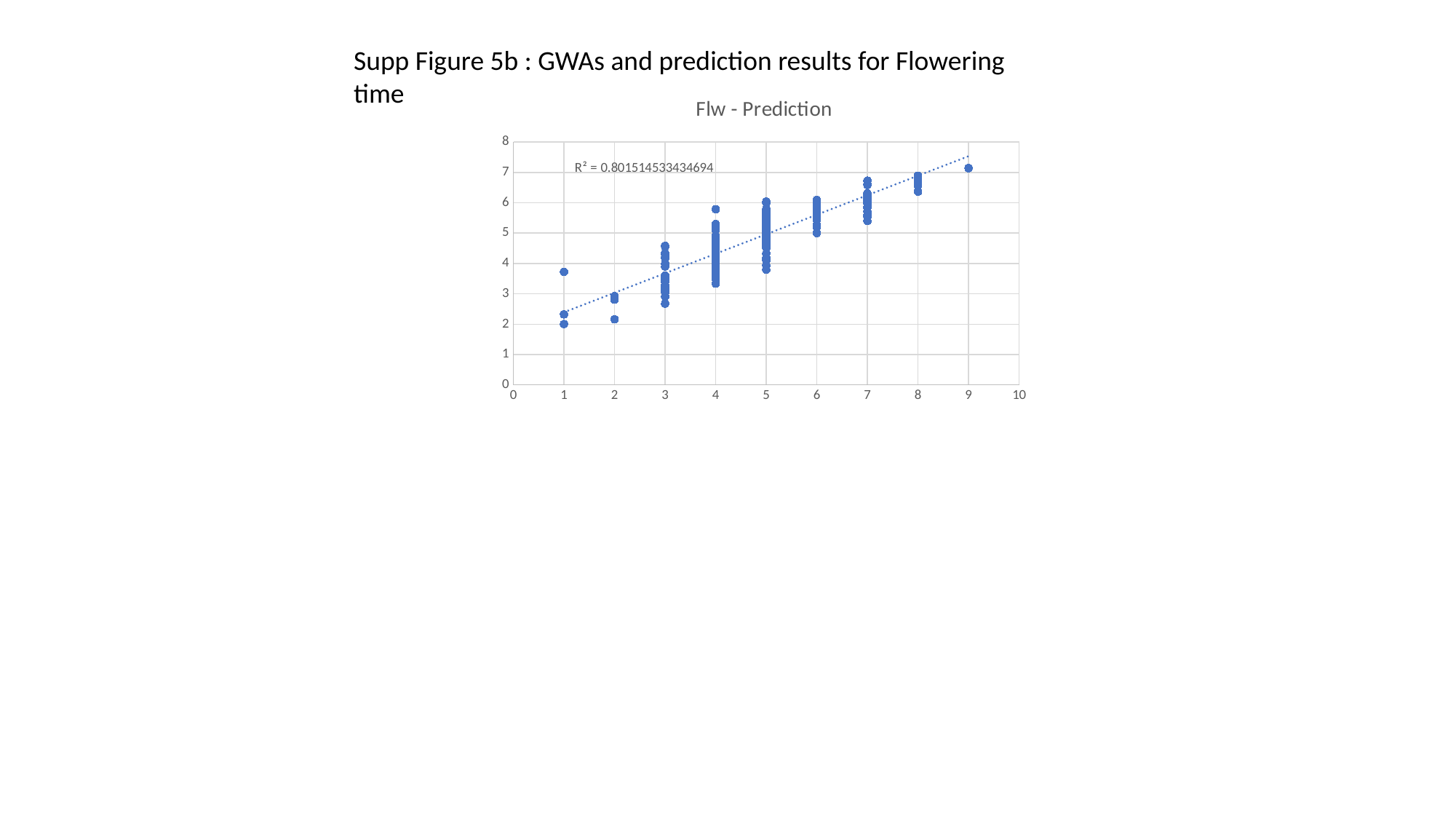

Supp Figure 5b : GWAs and prediction results for Flowering time
###### Chart: Flw - Prediction
| Category | Prediction |
|---|---|

### Slide 7
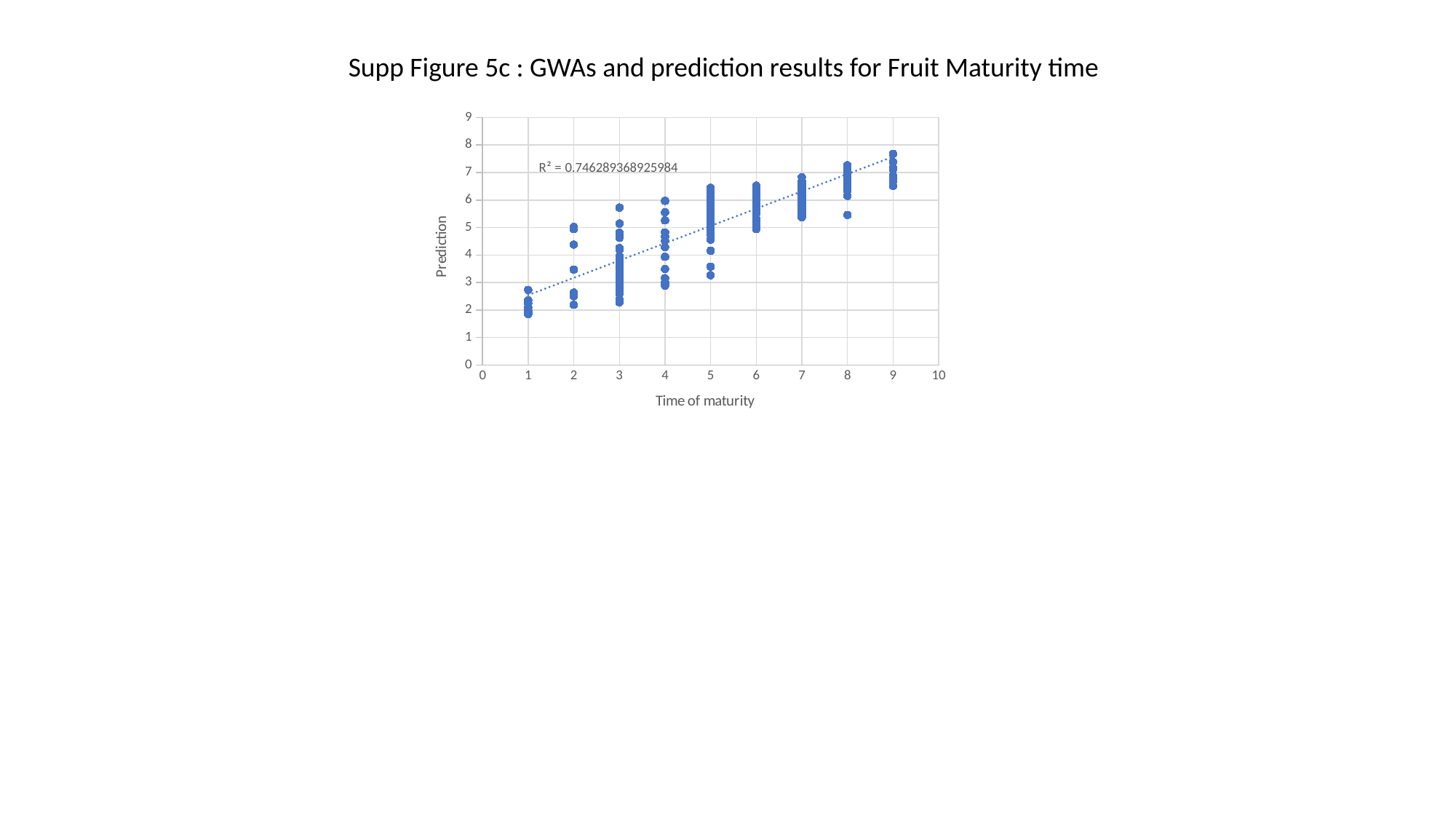

Supp Figure 5c : GWAs and prediction results for Fruit Maturity time
###### Chart
| Category | Prediction |
|---|---|
